# Quantitative spatial analysis of bacterial transcriptome and chromosome structural data with GRATIOSA: application to twin-supercoiled domain distribution

**DOI:** 10.1101/2023.12.22.573166

**Authors:** Maïwenn Pineau, Raphaël Forquet, Sylvie Reverchon, William Nasser, Florence Hommais, Sam Meyer

## Abstract

While classical models of transcriptional regulation focus on transcription factors binding at promoters, gene expression is also influenced by chromosome organization. Understanding this spatial regulation strongly benefits from integrated and quantitative spatial analyses of genome-scale data such as RNA-Seq and ChIP-Seq. We introduce Genome Regulation Analysis Tool Incorporating Organization and Spatial Architecture (GRATIOSA), a Python package making such combined analyses more automatic, systematic and reproducible. While current software focuses on initial analysis steps (read mapping and counting), GRAsTIOSA proposes an integrated framework for subsequent analyses, providing a broad range of spatially-resolved quantitative data comparisons and representations. As an example, we quantitatively assess the validity and extension of the twin-supercoiled domain model in *Escherichia coli* genome-wide transcription, using recent topoisomerase ChIP-Seq data. We show that topoisomerases are locally recruited by the 40% most highly expressed transcription units, with a magnitude correlating with the expression level. The recruitment of topoisomerase I extends to around 10 kb upstream, whereas DNA gyrase is recruited at least 30 kb downstream of transcription units. This organization is the primary determinant of topoisomerase I recruitment, whereas gyrase binding is additionally modulated at larger 100-200 kb length-scale. Further analyses of spatial regulation will be facilitated by GRATIOSA.

## Introduction

In bacterial cells, the chromosome is highly compacted through various factors, including DNA supercoiling (SC) and Nucleoid-Associated Proteins (NAPs), which together organize its structure dynamically through looping, bending, and twisting of the DNA (1, 2). In turn, the conformation of the chromosome affects DNA transactions, in particular transcription. The physical arrangement of DNA within the cell thus represents an “analog” information source crucial for the regulation of gene expression, which complements the “digital” information encoded by the genetic sequence (3, 4), but it largely escapes current modeling schemes.

High-throughput data such as RNA-seq, ChIP-seq and Hi-C, when combined, provide invaluable new information on this relationship (5–7). However, the analysis methodology should then differ from that inherited from classical, “digital” regulatory models based on transcription factors (TFs). In the latter, a regulator controls a gene depending on its sole promoter (“regulon”), independently from its genomic position and distance to neighboring genes. Software solutions currently used in the analysis of differential expression (8) and inference of regulatory networks (9) are all inspired by classical regulatory networks. Recently, combined software solutions were proposed for RNA-Seq and ChIP-Seq data, such as GenPipes (10), pyrpipe (11) and SnakeChunks (12) which facilitate the analysis of such data within a unified framework (downloading of raw sequencing reads, alignment, differential expression, peak calling, classification and Gene Ontology enrichment tests), but also rely on similar methodologies, and are therefore irrelevant to “analog” regulation. For example, numerous whole-genome data suggest that neighboring genes are co-expressed, even when they are unrelated in classical transcriptional regulatory networks (13), presumably because they belong to the same topological domain, and this feature is evolutionarily conserved (14). Analyzing this spatial regulatory interaction obviously requires considering the position of each gene, which none of the previously mentioned software does. In turn, the chromosome conformation is organized by abundant proteins (NAPs, topoisomerases) that bind the genome extensively and with looser sequence specificity than TFs, requiring a quantitative (more/less) rather than qualitative (bound or unbound promoter) analysis.

New analysis approaches are therefore necessary, for which no software solutions are available. Studies addressing these questions currently rely on homemade numerical analysis scripts, (14–17), with two major drawbacks: (1) specific skills and significant efforts are required for importing and comparing data of various types and formats, (2) reproducibility and standardization are more difficult to ensure. Here, we address this issue by presenting Genome Regulation Analysis Tool Incorporating Organization and Spatial Architecture (GRATIOSA), a new Python package targeted to computational biologists, and intended to facilitate such quantitative and spatial analyses of expression/structural data. It provides a unified framework for the automatic import and combination of many usual data types, and subsequent quantitative and spatial analyses of these data along bacterial genomes. Here “spatial” refers to the linear organization (along the genome sequence), where each gene occupies a well-defined position; the extension to 3D localization in the cell is addressed later in the manuscript.

GRATIOSA does not replace usual RNA-Seq or ChIP-Seq pre-analysis tools (read quality check, mapping, differential expression or peak calling, possibly using the integrated packages mentioned above), but rather facilitates the subsequent steps which previously required manual processing, i.e., the integration, comparison and combined statistical analyses of these data. It takes advantage of the versatility of Python for data analysis and numerical processing thanks to numerous available libraries. To the best of our knowledge, no comparable packages were proposed before as open tools for the community. In this article, we present the structure of the software, and illustrate its features through a specific but important example of “analog” regulation: the quantitative interplay between gene expression and topoisomerase activity along the *Escherichia coli* genome, which has never been analyzed in a global and systematic way.

## Materials and Methods

### Framework

GRATIOSA is written in several Python source files, each defining one of the main object classes (Genome.py, Gene.py, GO.py, Transcriptome.py, Chipseq.py, …) with their attributes (e.g., a Genome object contains Genes and a genomic sequence) and methods (e.g., load_annotation, load_seq, …). In addition, some miscellaneous functions for each class are defined in a separate source file (e.g., intermediate functions for reading a gff file are defined in useful_functions_genome.py). Finally, three additional program files contain intermediate functions for statistical tests (stat_analysis.py), and the graphical representation of genomic data (plot_genome.py) or of statistical results (plot_stat_analysis.py). All methods and functions rely on the packages NumPy (18), Matplotlib (19), pandas (20), pysam, SciPy, and Statsmodels. Three main types of interdependent objects can be distinguished: the Genome object, the elementary objects (Gene, TU, TSS, TTS objects), and the objects containing experimental data (Transcriptome, ChIP-Seq, Hi-C objects) (Supplementary Fig. S1). An elementary object represents a single gene or a single TU (Transcription Unit), while the Genome object contains the annotation of all elementary objects (i.e., all genes, all TUs). Similarly, the elementary object may have attributes such as expression level or average ChIP-Seq signal for its segment, but the overall expression levels and genomic signals are attributes of the Transcriptome and ChIP-Seq objects, respectively.

### Experimental data pre-processing

GRATIOSA can import experimental data formatted either as continuous signals (i.e., one value per genomic position), discrete sites (i.e., list of genomic positions), or gene-wise values (i.e., one value per gene). The data files follow usual formats generated with standard external tools such as DESeq2, MACS, or deepTools (Table 1). Functional enrichment analysis using Gene Ontology (GO) terms is also performed using standard description files, namely an annotation file for the analyzed species and a file describing all GO terms (obo, available from the GO website (21)). The annotation file (typically generated by software such as Blast2GO) contains at least two columns, with each row relating one gene with one GO (with multiple lines for each gene). The obo file can be automatically downloaded with the package.

**Table 1.** Summary table of the data types analyzed by the package (mostly tabular files), with their content information (rows and columns) and typical software used to obtain them. The last row describes the import of a wide range of datasets describing lists of discrete genomic sites (e.g., Hi-C domain borders, experimental or predicted protein binding sites, list of other genomic features, …).

| Data type | Rows | Columns | Standard format | Commonly used packages |
| --- | --- | --- | --- | --- |
| RNA-Seq coverage | positions | coverage | bedgraph | Bedtools (22) |
| RNA-Seq expression | genes | locus, expression level (RPKM or FPKM) | tab | Cufflinks, HTSeq (23) |
| RNA-Seq differential expression | genes | locus, fold-change, p-value | csv | limma (24), DESeq2 (8) |
| Microarray differential expression | genes | locus, fold-change, p-value | csv | limma (24) |
| ChIP-Seq coverage | positions | bin start, bin end, coverage | bedgraph | deepTools (25) |
| ChIP-Seq peaks | sites | position, value | bed | MACS (26) |
| list of genomic position data | sites | position, value | csv |  |

### Database organization

To be analyzed with GRATIOSA, data must be organized in a database with a fixed structure: each directory represents one organism (with a reference genome) and contains multiple subdirectories (Supplementary Fig. S2). The annotation subdirectory contains the annotation, usually in gff3 format. All other subdirectories have the same structure, each containing experimental data files and an info file (See Experimental data pre-processing). chipseq and hic subdirectories are further subdivided, to distinguish between two types of data (peaks and signals in the case of ChIP-Seq, loops and borders in the case of Hi-C), each following the same structure as other subdirectories.

### Data import

Data import is automatic from the annotation files (sequence, annotation). For experimental data present in multiple files (e.g., different experimental conditions for gene expression), the import function uses an “info” file, i.e., a tab-separated text file containing the list of data files that will be loaded for analysis (Supplementary Fig. S3). This method enables the automatic selection and import of data from various sources or softwares, whether in standard or custom formats, by specifying some descriptors of each data file. For example, in the case of the expression directory describing genome-wide expression levels in various conditions (microarray intensities or typically for RNA-Seq data, RPKM/FPKM values), the info file contains a list of rows (one per experimental condition to be loaded), each indicating the condition name, associated file name, column index for gene names/tags, column index for expression data, whether the data is in log format or not, and the separator used in the file.

Depending on the type of data to be analyzed, the corresponding object needs to be initialized first (here Transcriptome), and then data import is achieved using the appropriate method (e.g., load_expression to import expression data onto the Transcriptome object, see Supplementary Fig. S3). Intermediate functions (e.g., useful_function_Genome.read_seq) are used to read data files, and might be called by the user in unusual cases (e.g., files with very custom formats).

### Data processing

Once data are loaded (typically as Python lists, NumPy arrays, or Pandas dataframes), new attributes can be either added manually or computed from existing ones. For example, ChIP-Seq signals can be binned, smoothed, or averaged. For binning, the user defines a bin size and method (by default, average signal within the bin); for smoothing, the size of the sliding window; for averaging, the list of different signals to be used (usually replicates). Raw or computed signals can be automatically related to Gene objects by computing the average signal over the gene region. Similarly, various attributes (such as orientation, expression level or fold-change, protein occupancy) can be added to Gene objects based on imported Genome, Transcriptome or Chipseq attributes (Supplementary Fig. S1, attributes written in italics). An orientation can be defined for a gene or intergenic position, depending on the direction of neighboring genes (tandem, divergent, convergent). Before performing statistical analyses, quantitative data can be classified into qualitative categories, based on user-defined thresholds or on quantiles (of equal or custom sizes).

### Computational resources

GRATIOSA is designed to run on a standard desktop/laptop computer, coming after the most computationally intensive steps in sequencing data analysis (especially read mapping). Usual operations take less than a few seconds with < 1 Gb memory load. Most data types (annotation, gene-wise expression values) have a low (a few Mb) memory load. Data types relying on genomic coverage (RNA-Seq, ChIP-Seq) are usually larger but, because of the small size of bacterial genomes, are also easily tractable (dozens of Mb; optionally, input tabular files can be automatically converted into binary NumPy array files that are lighter and faster to load).

### Statistical analysis and outputs

The package offers three standard statistical tests (Supplementary Table S1): an enrichment test (hypergeometric, with false discovery rate correction), particularly useful for functional enrichment analysis; a proportion test (based on the z-score, equivalent to a chi^2^ test), for example for comparing the proportion of differentially expressed genes in different sets of genes; and a standard Student’s t-test for quantitative data, for example for comparing the average expression level or fold-change among different sets of genes. These tests are implemented using common Python modules, and their results are saved in a csv file.

### Graphical functions

Statistical results can be visualized as bar plots generated using the matplotlib.pyplot.bar module, with error bars representing 95% confidence intervals, and optionally, an annotation of these bar plots with brackets and stars according to their significance level (* for p-value < 0.05, ** for p-value < 0.01, and *** for p-value < 0.001). Other graphical functions are implemented in the package to visualize signals, such as ChIP-Seq and RNA-Seq coverages within their genomic context (Fig. 1). RNA-Seq coverage corresponds to the number of reads for each genomic position, on each strand. Depending on the size of the visualization window (from a few genes to the entire genomes), genes on each strand can be annotated.

**Fig. 1.**
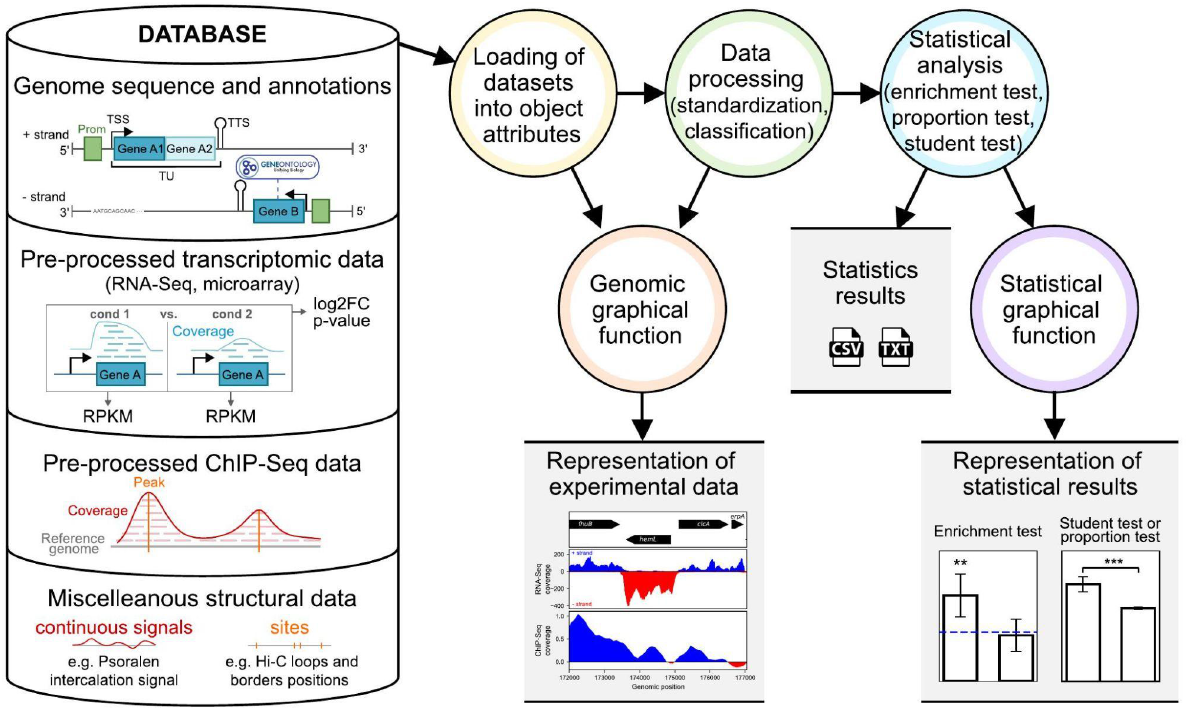
Graphical overview of the GRATIOSA package. The left section (database) presents the different types of data processed by the package. The right section shows the main steps and outputs of the package. For a more detailed description, see Supplementary Figure S3.

### Processing of *Escherichia coli* topoisomerase and expression signals

The Escherichia coli W3110 topoisomerase signals analyzed in this article were previously published by Sutormin et al., with FPKM values of annotated transcription units (TUs) (5, 16). The mapping of gyrase cleavage sites was obtained using a method called Topo-Seq; we used the datasets obtained with ciprofloxacin as gyrase poison (GEO: GSE117186) (5). Topoisomerase I binding sites were identified with ChIP-Seq (GEO: GSE181915) (16), with or without rifampicin treatment (an inhibitor of transcription). For each sample, the coverage was first normalized by its mean, and the log2 ratios between immunoprecipitation (IP) and control (mock) coverage were computed. The log2 ratios were then normalized (by their mean) and the resulting values were saved in bedgraph format compatible with GRATIOSA. The replicates of topoI and gyrase signals were averaged for each genomic position (using the Chipseq.load_signals_average method), representing the log2 occupancy by each protein relative to the control data. The signals were then separated into macro-scale and micro-scale contributions (see Results): the macro-scale contribution was obtained by smoothing each signal along 200-kb sliding windows (with useful_functions_Chipseq.smoothing); the micro-scale contribution was obtained by subtracting the macro-scale contribution from the raw signal. For expression analyses, we generated approximate expression signals (per genomic position) from the FPKM (per TU) expression values provided (16). Pearson correlation coefficients for signal autocorrelation were calculated using the scipy.stats.pearsonr function by comparing each signal with the same signal shifted in genomic position. Correlation coefficients were computed using the same SciPy function to topoisomerase signals and the expression signal, after the smoothing of these signals at different scales.

## Results

### The GRATIOSA package

GRATIOSA is written in Python and runs on a standard desktop or laptop computer. It facilitates the integration and comparison of bacterial genomic data types using standardized statistical analyses, and is particularly designed to investigate quantitatively the spatial relationship between transcriptional regulation and chromosome organization. Two basic types of experimental data can be analyzed with the package: expression data (transcriptomics) and structural data (ChIP-Seq or related technologies), and both can be represented either in a continuous (RNA-Seq or ChIP-Seq coverage) or in a discrete manner (gene expression levels, positions of ChIP-Seq peaks). An overview of the package is shown in Fig. 1, and more detailed technical information is given in Materials and Methods. The simplest method to import data with GRATIOSA is to organize the data files (in usual formats) following a predefined structure, classified by organism and data type (Supplementary Fig. S2). They can be accompanied by an information file enabling an automatic selection and import of these data (e.g., a subset of experimental conditions for a particular analysis of expression data). This basic structure can be used to import many other and/or custom types of data, e.g., predicted protein binding (discrete) sites or (continuous) signals, miscellaneous annotated features, positions of Hi-C domain borders, etc.

The GRATIOSA package is typically imported into a Jupyter notebook, and data analysis is executed as Python commands, in three major steps (Fig. 1). First, different objects (Genome, Transcriptome, Chipseq, etc; see Supplementary Fig. S3, gray) are initialized, depending on the user’s needs. The required data are then loaded as object attributes (Supplementary Figs. S1 and S3, yellow).

During this second step, the loaded attributes are processed by the software (Fig. 1) and become easily manipulable (mainly as NumPy arrays or Pandas dataframes), either manually (for custom computations) or using the tools implemented in the package (described in Supplementary Figure S3, green). Among the latter, attributes associated with genomic positions (“signals”) can be scaled to genes (for example, the average ChIP-Seq coverage is computed for each gene). Signals can be smoothed. Genes can be classified according to their orientation (with respect to neighbors) or length. Thanks to the easily manipulable formats, the user can perform custom additional processing to modify or add attributes, possibly using manually imported external data. A verification and graphical exploration of the signals can be performed at the end of the first or second steps using a graphical function that automatically plots signals with the annotated genome. Finally, to prepare for statistical analysis, quantitative data (expression level, protein binding strength, etc.) can also be classified using thresholds or class sizes, e.g., in expression deciles.

The last step is the statistical analysis (Fig. 1, Supplementary Fig. S3, blue) with enrichment or proportion tests (for qualitative attributes) or Student’s t-tests (for quantitative comparisons), as described in Materials and Methods. Test results are exported as tables (in csv format) and can be visualized as annotated bar plots created with graphical functions included in the package (Supplementary Fig. S3, purple).

These steps are illustrated in the following case study, for which GRATIOSA allowed a drastic reduction in code writing and improvement in reproducibility. The associated codes are given as Jupyter notebooks in Supplementary Files 1 and 2, and have been deposited in GitHub, as well as tutorials illustrating a few typical analyses.

### Characterization of *Escherichia coli* twin-supercoiled domains *in vivo*

DNA supercoiling (SC), i.e., torsion-induced deformation of the DNA double-helix, besides its essential role in bacterial chromosome compaction, is recognized as a global transcriptional regulator (27–29). The superhelical level is controlled globally by the action of topoisomerases, mostly topoisomerase I (topoI) and gyrase (30), but is also strongly affected by local transcription, according to the “twin-supercoiled domain” (31), leading to the accumulation of negative (resp. positive) supercoils behind (resp. ahead of) the elongation complex. However, until recently, there was limited information on twin-domains in the chromosome.

Since topoI preferentially binds to negatively supercoiled regions (where DNA forms transient denaturation bubbles), and DNA gyrase to positive supercoiled regions (forming plectonemes), mapping the activity of these two enzymes provides spatially resolved information on the supercoiling distribution. Recently, Sutormin et *al*. used ChIP-seq of topoisomerase I, and a ChIP-Seq derived assay mapping cleavage intermediates of gyrase (5, 16), to analyze the genome-wide binding (resp. cleavage) distributions of these two enzymes in *E. coli* W3110, thus providing invaluable insights into the presence, amplitude, and extension of twin-supercoiled domains *in vivo*. These authors found enrichments of topoI upstream and of gyrase downstream of the 200 most highly expressed transcription units (HETUs) out of approximately 2100 annotated TUs in *E. coli* (16), in agreement with the twin-supercoiled model. We used GRATIOSA to characterize these twin-domains more systematically and quantitatively. Is this behavior specific to that small minority of genes? How many genes exhibit a local enrichment in topoisomerase recruitment? To what distance does it extend? Can it be mapped to the formation of topological domains?

We started from the raw data and computed coverage distributions for the different samples using standard methods (see Materials and Methods). All data import and most classification steps could then be achieved with few lines of code with GRATIOSA, allowing the user to focus on the specific analyses and plots shown in the following (Supplementary Files 1 and 2).

### Distribution of supercoils around transcription units of various expression levels

We first wished to know how many genes were associated with detectable twin-domains. We classified all TUs by expression level and analyzed the average gyrase and topoI coverages (using log2-normalized ratios of IP-signals versus mock, with an average level of 0) from 10 kb before TU start to 10 kb after TU end (Fig. 2).

**Fig. 2.**
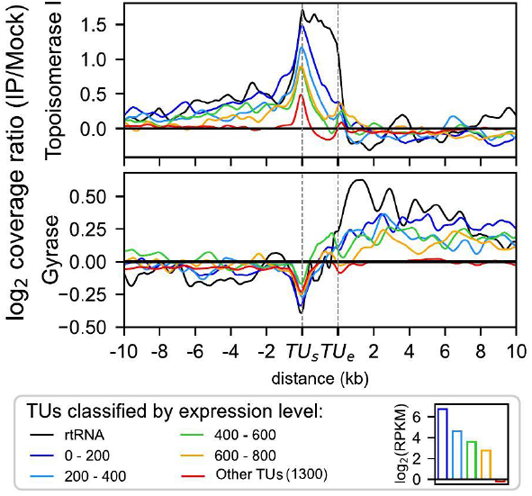
TopoI binding (top) and gyrase cleavage (bottom) signals around TUs in *E. coli* (ChIP-Seq data from (5, 16)). Shown signals are normalized log2 ratios between coverage after immunoprecipitation (IP) and control coverage (mock). TUs were classified based on their expression level, with 200 TUs in each class of highly expressed TUs (around one decile each: blue, cyan, geen, orange), one class with all other protein-coding TUs (red), and one class with strongly expressed TUs encoding stable RNAs (ribosomal or transfer RNAs, black). By construction, both signals have a genomic average of 0. The bottom right panel shows the average expression level of each expression class, except for stable RNAs that have the strongest expression (but for which FPKM do not reflect the expression level). TU_s_ and TU_e_ indicate the start and end of the TU.

The topoI binding signal (Fig. 2, top) exhibits a clear gradual increase over 10 kb upstream of the TU start, with a pronounced peak at the TU start. Following the TU start, the signal quickly decreases and becomes close to zero by the end of the TU (except for stable RNAs in black). Within the TU, higher expression levels are associated with higher topoI binding density, consistent with the known colocalization and interactions between topoI and RNAP (16, 32, 33). Within stable RNA (tRNA and rRNA) operons (black curve), topoI binding appears relatively flat, potentially indicating an ongoing topoI recruitment throughout the TU, in contrast to all other categories (even strongly expressed). This observation suggests a specific extensive requirement for topoI for those operons. Upstream of the transcription start site, the density of topoisomerase I binding is also markedly dependent on TU expression levels. This enrichment is likely not due to the interaction of topoI with RNA Polymerase, but reflects the negative supercoils generated during elongation, in clear agreement with the twin-supercoiled domain model (31). *In vivo*, based on the ChIP-Seq data, this phenomenon affects not only 200 HETUs, but at least the 800 most highly expressed TUs (approximately 40% of the genome). It extends remarkably far: up to 10 kb upstream of the TU for the top 400 most highly expressed TUs, up to 8 kb for the next 200 TUs, and up to 6 kb for the following 200 TUs. Thus, the apparent extension of the twin-supercoiled domain is dependent on the TU expression level. In the 1300 (∼60%) weakly expressed TUs (red), no enrichment is observed. After the end of the TU (downstream), the topoI binding is slightly weaker than the genomic average (negative levels for all expression classes), as would be expected from the twin-domain model if positive supercoils inhibit topoI binding.

The gyrase signal (Fig. 2, bottom) exhibits significant differences compared to that of topoI. As expected again, the 10 kb upstream of TUs are less cleaved by gyrase than the genomic average, possibly due to the negative supercoils generated by transcription, with a particularly strong depletion at the beginning of the TU. Within the TU, we observe no enrichment in gyrase binding, suggesting that the enzymes are not required in the immediate vicinity of RNAP (in contrast to topoI). After the TU, the signal increases sharply, then slowly decreases but remains strongly enriched after 10 kb behind the TU, in all classes of the 40% highly expressed TUs, and thus extends further and more uniformly than the topoI enrichment before the TU. This enrichment is clearly increasing with expression strength, whereas the 60% weakly expressed TUs exhibit no enrichment.

We wish to emphasize that, because of the high density of bacterial genomes, the 10 kb regions flanking the central TU contain many other genes (one TU every 2 kb in average, with short intergenic regions). The expression of the central TU and of these neighbors may both contribute to the observed topoisomerase binding patterns. Even if the obvious regularity of these patterns suggests a strong impact of the central one under investigation (whereas the neighboring TUs differ by their positions, expression strengths, and orientations), it does not constitute evidence that supercoils actually propagate over more than 10 kb from the center, especially if the neighbors’ expression is correlated to the central TU’s and co-directionally oriented. Further correlation analyses are developed in the upcoming paragraphs, and their interpretation is addressed in the Discussion.

### Role of gene expression strength and orientation in shaping local supercoiling distributions

We now further investigate the correlation between expression level and twin-supercoiled domain intensity, by examining the total binding signals around all TUs separated in expression deciles, in the 10 kb upstream of the TUs (Fig. 3, left panel), within the TUs (Fig. 3, middle panel, expression levels are indicated in gray), and in the 10 kb downstream (Fig. 3, right panel).

**Fig. 3.**
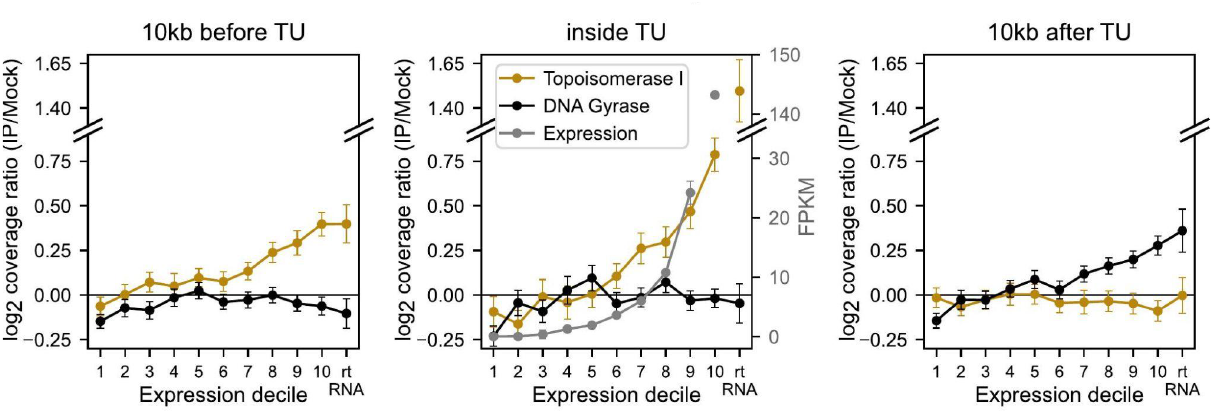
Average topoI binding (gold) and gyrase cleavage (black) signals in the 10 kb before (left), inside (middle), and in the 10 kb after (right) TUs, depending on the expression level. TUs are classified in expression deciles, and the average expression level (FPKM) of each decile is shown in gray in the middle panel; operons of stable RNAs are classified separately. Mind the breaks in the vertical axes. Error bars indicate 95% confidence intervals.

This representation confirms the previous observations, with the following additional information. Within TUs, the average density of topoI (Fig. 3, gold curve middle panel) exhibits a moderate increase in the first deciles of TU expression levels, which becomes much more pronounced in the last 4 deciles (i.e., the 40% most highly expressed TUs). In the 10 kb region upstream of the TUs (Fig. 3, gold curve left panel), the expression level dependence is very low in the six first deciles, whereas topoI binding steadily increases in the last four deciles. After the TUs (Fig. 3, gold curve right panel), as observed in Fig. 2, the topoI binding signal is weakly negative for all expression levels, suggesting a weak accumulation of positive supercoils.

As for gyrase (Fig. 3, black curves), its recruitment before and within TUs is relatively constant, even in the most expressed ones. Before the TUs, the signal is slightly negative, compatible with the presence of negative supercoils repelling gyrase. After the TUs, a gradual enrichment of gyrase cleavage with higher expression is observed, especially once again in the last 4 deciles, and quite limited before.

Together with those of Fig. 2, these results (1) systematically confirm the *in vivo* relevance of the twin-supercoiled domain model; (2) consistently suggest that topological constraints affect not only a small minority, but the 40% most highly expressed TUs, with a clear and steady increase in the recruitment of topoisomerases depending on the expression level. The 60% remaining weaker TUs are less affected, presumably because they generate moderate mechanical distortions that might be (at least partly) dissipated without local recruitment of topoisomerases; (3) within TUs, topoI is recruited in the wake of the elongating RNA Polymerase (16), especially close to the promoter, but gyrase is not enriched.

Previous studies have shown that, because of the asymmetry of transcription-induced supercoils, local gene orientation is important for the transcription-SC coupling (15, 16, 34, 35). The package has built-in functions to classify intergenic regions depending on the orientation of their neighbors (divergent, convergent, tandem), and quantitatively compare how they recruit topoisomerases. We observed (Fig. 4) an enrichment of topoI and a depletion of gyrase between divergent genes compared to convergent genes, as already noted by Sutormin et *al*., following expectations of the twin-supercoiled domain model (16). By comparing the signals from these regions with and without rifampicin, an inhibitor of transcription, we observed that this difference was still present, to a lesser extent, after the treatment (Fig. 4 B and E). This effect can be explained by differences in base sequence (especially since divergent regions are more enriched in AT-rich promoters, Supplementary Fig. S4), i.e., binding preferences independent of transcriptional activity. It might also reflect an incomplete dissipation of supercoils generated by previous transcriptional activity (before rifampicin treatment) or an incomplete transcriptional inhibition by the drug. By normalizing the untreated coverage (Fig. 4 A and D) with the coverage after rifampicin treatment (Fig. 4 B and E), we erase out any static effect, and confirm a differential and opposite recruitment of topoisomerases, specifically induced by transcriptional activity depending on the local gene orientation (Fig. 4 C and F). For topoI, the enrichment is symmetrical, with div regions exhibiting enrichment and conv regions exhibiting depletion, while tandem regions are close to 0 (i.e., the genomic average). In contrast, for gyrase, transcription induces no specific enrichment in convergent regions (close to genomic average), whereas tandem and div regions are depleted. This difference suggests that transcription of convergent genes does not lead to a significant buildup of positive supercoils and subsequent recruitment of gyrase between them, whereas the negative supercoils upstream of TUs efficiently reduces its binding in tandem and especially divergent regions.

**Fig. 4.**
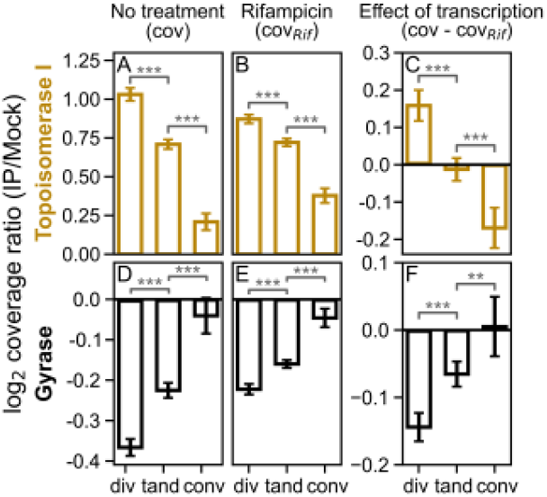
TopoI binding sites (gold) and gyrase cleavage sites (black) density in intergenic regions, depending on the orientation of surrounding genes. Each bar represents the average normalized log2 ratios between coverage after immunoprecipitation (IP) and control coverage (mock) for a specific orientation, calculated after binning intergenic positions into 100 bp bins. Curves in **A** and **D** were obtained without treatment, **B** and **E** after treatment with rifampicin, a transcription inhibitor, and **C** and **F** show the difference between untreated and rifampicin-treated samples. Error bars are 95% confidence intervals.

### Extension of twin-domains and multiscale interplay between transcription and topology

Fig. 2 showed that topoisomerase recruitment was affected at least up to 10 kb before and after TUs, close to the most commonly accepted size for topological domains in *E. coli* (36). We now look at the extension of topoisomerase binding correlations possibly reflecting these domains. The binding autocorrelation curves (computed along the whole genome) are shown in Fig. 5A. Unsurprisingly, the correlation is strong below one kilobase (the typical gene length), for both topoisomerases and for gene expression patterns, as expected if short-scale binding variations are related to transcription (and close to the spatial resolution of ChIP-Seq data). The correlation then reduces with distance, but in a very different manner for the two enzymes: the characteristic length-scale of the topoI binding distribution is 10 kb, above which it is entirely decorrelated; in contrast, at that scale, the correlation of gyrase binding is still quite strong, and vanishes only a full order of magnitude above, after 100 kb. These two separate length-scales were already described in earlier works, and correspond to two different levels of genome organization: (1) the 10-kb (hereafter quoted “micro”) scale is that of transcription units and their immediate neighboring interactions, possibly within transient topological domains (36), leading to strong and evolutionarily conserved co-expression (14); (2) the larger 100-200 kb scale (“macro”), already observed in transcription and gyrase binding signals in *E. coli* (13) as well as in 3D conformation maps (1, 37). The curve of gene expression (gray) confirms this separation, with the correlation dropping almost entirely at the micro-scale, but a small (∼10%) contribution still observed up to the macro-scale. Accordingly, the correlation of topoisomerase binding with expression is maximal at length-scales of a few dozen kb (topoI) vs 200 kb (gyrase).

**Fig. 5.**
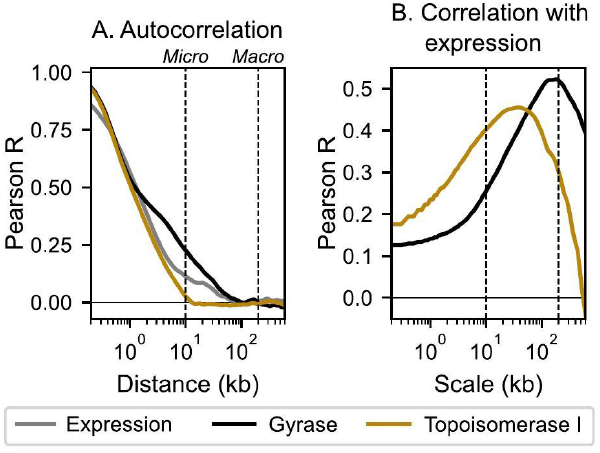
**(A)** Pearson coefficients for the autocorrelation of gyrase cleavage (in black), topoI binding (gold), or expression signal (gray) (as a function of distance). **(B)** Pearson coefficients for the correlation between gyrase (black) or topoI (gold) and expression signal, smoothed at increasing scales. Similar curves are obtained with Spearman’s coefficient (not shown).

This observation suggests that topoisomerases are recruited based on two (partly) separate contributions in terms of mechanisms and length-scale: (1) micro-scale transcription and (2) macro-scale 3D structuring of the chromosome. To test this hypothesis further, in Fig. 6, we used a signal filtering method to compute the contributions of these two length-scales in the variations in binding distributions (by smoothing over the macro-scale of 200-kb, vs subtracting the smoothed signal and retaining only the micro-scale variations, see Materials and Methods). At the micro-scale, as expected from Figs. 2-3, both enzymes exhibit a similar amplitude of binding variations (vertical standard deviations) directly related to local expression (the colored dashes follow the expression strength), suggesting that transcription is indeed the major driver of topoisomerase recruitment at that scale. In contrast, the two enzymes behave differently at the macro-scale. Gyrase binding differs significantly among the different macro-regions of the chromosome (vertical standard deviation), with the same range of variability as that observed at the micro-scale; in contrast, the recruitment of topoI is relatively uniform at the macro-scale, showing that micro-scale transcription alone, rather than the large-scale organization of the chromosome, determines the recruitment of that enzyme.

**Fig. 6.**
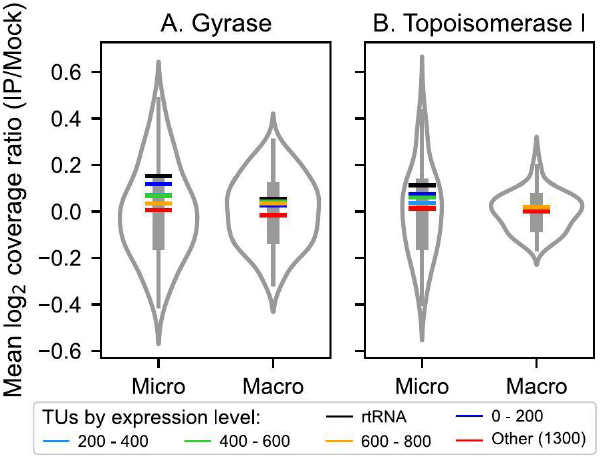
Distributions of **(A**) gyrase cleavage and **(B)** topoisomerase I binding magnitudes across macro-scale regions of the chromosome (200 kb, corresponding to large-scale 3D structuring of the chromosome) versus micro-scale variations (10 kb, dominated by transcription) within these regions. The contributions were computed by signal filtering (see Materials and Methods) and binned at 50 kb. Colored lines indicate the average signal after (gyrase) or before (topoI) different classes of TUs. Colors: rtRNA (black), 200 most highly expressed TUs (dark blue), and following classes of 200 TUs by decreasing expression (cyan, green, orange), and 1300 remaining TUs (red).

Focusing now on transcription at the micro-scale, to what distance does the expression of a specific TU affect topoisomerase recruitment? We addressed this question in Fig. 7, by computing the cumulative binding (area under the binding profile) between the TU start/end and a remote position up to a distance of 50 kb: a flat curve thus indicates a local binding intensity similar to the genomic average (and thus presumably no influence of the TUs under consideration), whereas an increase (resp. decrease) indicates a stronger (resp. weaker) binding, possibly due to transcription.

**Fig. 7.**
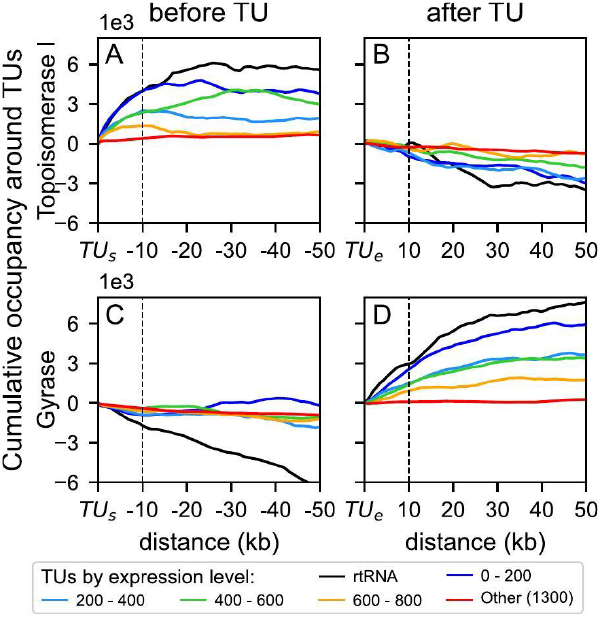
Extension of the transcriptional-induced topological constraints. The top and bottom panels show cumulative signals (i.e., total binding from the TU) for TopoI binding and gyrase cleavage, respectively, in the 50 kb before the TU start (left panels) and after the TU end (right panels). TUs were classified based on their expression level (same color legend as in Fig. 2). The plotted signals correspond to the average area under the curve of the micro-scale signal (i.e., after subtracting the macro-scale contribution, especially for gyrase), from the TU start or end site to increasing distances (up to 50 kb).

As already observed, topoI binding increases with the expression level in the first 10 kb before the TU, and the extension of the transcription-induced negative supercoils also tends to increase (Fig. 7A). For stable RNAs (black curves), the signal reaches 25-30 kb, consistent with the size identified around the ribosomal operons by psoralen binding (6). After the TU (Fig. 7B), the depletion in topoI binding (negative values) mostly occurs in the first 10-20 kb, and slightly increases with the expression level. For gyrase, no specific signal is detected upstream of the TU start (C), except for stable RNAs, where the enzyme appears to be repelled over at least 50 kb, possibly due to strong negative supercoils accumulating there. Downstream of the TU (Fig. 7D), a strong enrichment in gyrase cleavage is observed, that increases with the expression level; but in contrast to topoI, it extends way beyond the 10 kb region, from around 20 kb for moderately expressed TUs (orange) to around 40 kb for strongly expressed ones (dark blue) and even beyond for stable RNAs. Thus, positive supercoils induced by transcription may extend as far as several dozen kilobases. We checked that neighboring TUs exhibit similar expression levels regardless of the class of the central one (Supplementary Fig. S5), suggesting that the latter could be the main driver of these far-reaching supercoils, as developed further in the Discussion.

## Discussion

### A software solution for integrative spatial analysis of genomic data

The interplay between chromosome conformation and gene expression is a complex and intricate problem, requiring dedicated conceptual and technical tools. While classical regulatory networks are usually analyzed in a qualitative way (does a regulatory protein bind at a promoter?), the “analog” layer of transcriptional regulation can be only understood through quantitative analysis (does it bind more or less?), considering spatial (1D or 3D) relations between genes, and combining different actors (architectural proteins, SC, expression, …). GRATIOSA facilitates such analyses, by automating the formatting and import of various data into a uniform framework, providing a range of standard tests and visualization tools, and facilitating quantitative analyses of different data types with various Python tools and libraries. This is illustrated in the example of topoisomerase distributions, where the basic steps of the analysis are standardized, and the programming effort can be dedicated to the design of specific statistical tests and graphs (see notebooks in Supplementary Files). While the original analyses of these data (5, 16) were restricted to a small minority of highly expressed genes exhibiting the strongest effects, GRATIOSA provided the ideal framework to extend these analyses and characterize for the first time the twin-domains *in vivo* in a systematic manner.

The present version of the code focuses on the spatial organization along the 1D genomic sequence (linear), based on well-defined gene positions, which is usually disregarded in regulatory analyses. A natural extension of the software is the inclusion of the 3D organization of the chromosome through Hi-C data, with substantial additional issues to consider: (1) the relation between Hi-C contact densities and 3D conformation (both static and dynamic) is nontrivial and relies on complex modeling; (2) the extension of the package to automated 3D analyses will involve significantly more complex functions. The present version of the software already offers limited solutions for Hi-C data, through a prior analysis by dedicated software (38) providing lists of genomic positions with specific identified features (e.g., borders between chromatin interaction domains, or loops) which can then be imported by GRATIOSA and treated as unidimensional features (e.g., in order to analyze the gene expression level and orientation, or protein binding enrichment, at the borders between Hi-C domains).

### A quantitative description of twin-supercoiled domains in the chromosome

We addressed how many genes exhibit local enrichment in topoisomerase recruitment, presumably reflecting the presence of strong topological constraints, and how far these constraints extend, both questions of significant importance but still largely unanswered.

The analyzed data consistently confirm the widespread relevance of the twin-domain model in bacterial genomes (Fig. 2). The phenomenon is detected specifically in the 40% highly expressed fraction of the genome, where it exhibits a clear and increasing magnitude with the expression strength (Fig. 3). For the 60% remaining genes, weaker supercoils might still exist but escape the sensitivity of the ChIP-Seq-type methodology; however, the absence of detectable signal suggests that the weak transcription level of more than half *E. coli* genes can be sustained without local recruitment of topoisomerases. Topological effects are strongly dependent on the local orientation of neighboring genes (Fig. 4), suggesting that topoI is not required at the average expression level at tandem genes, but is strongly recruited between divergent promoters, as observed long ago in *S. enterica* (39), in addition to its strong activity at highly expressed genes. In contrast, while gyrase is repelled from divergent regions by negative supercoils, it is not specifically attracted by the expression of convergent genes (Fig. 4F). This suggests that, at least in the analyzed experimental conditions, transcription is usually not hampered by accumulating positive supercoils, even in convergent genes. This observation is fully consistent with recent results showing that the expression of a single gene is dependent on the distance to a topological barrier on the upstream but not the downstream side (40), and that gyrase inhibition does not usually inhibit but surprisingly activate convergent genes (15). RNAP stalling because of positive supercoils is therefore maybe not an extensive phenomenon *in vivo*, but rather limited to a few highly expressed genes (at least in the exponential phase where the baseline superhelical level is strongly negative).

What is the extension of twin-supercoiled domains, and the relative magnitude of transcription vs other mechanisms in shaping the local distribution of supercoils? This question is important, as theoretical models suggested that the propagation of supercoils might induce collective expression modes between neighboring genes in the densely packed bacterial genomes (41–43). The spatial analysis (Figs. 5-6) shows that topoI recruitment is strongly dominated by transcription, with a characteristic length-scale of 10 kb; in contrast, gyrase contains two different contributions of comparable magnitude, one from transcription (at a comparable scale), and one an order of magnitude larger (100-200 kb). The former spatial scale is remarkably close to that previously proposed for topological domains, based on different data (36). The latter scale was already identified in early analyses of transcription and gyrase binding patterns (13), as well as 3D Hi-C maps (“chromatin interaction domains”) (1, 37). The underlying mechanisms remain ill-defined, but presumably implicate a complex interplay of architectural proteins, transcription and replication-related factors (1). Interestingly, a recent study also exhibited a statistical relation between Hi-C contact density and GapR binding (indicative of positive supercoils), but not topoI (indicative of negative supercoils) (17). Since GapR and gyrase are expected to bind similar positively supercoiled regions, that result is fully consistent with our observation of a predominant relation of gyrase binding with large-scale chromosome structure.

### Transcription and supercoiling: from genomic correlations to mechanistic causality

At the micro-scale, the separation of the 40% strongest TUs into expression classes clearly suggests that transcription is the dominant factor in the recruitment of gyrase and topoI, following a strong linear (rather than 3D) ordering (Fig. 7). TopoI is strongly recruited along transcription units (especially near the TSS), as expected from its interaction with RNAP (33), but also upstream of them, up to 10-20 kb, increasing with the expression level. Gyrase binds downstream of TUs up to 30-40 kb, not only at highly expressed operons but at almost half of them. The proposed analysis is, to our knowledge, the first to provide relatively direct evidence of such extended topological distributions related to transcription in a significant fraction of the genome.

Does this mean that supercoils generated by transcription (of the central TU in our expression classes) actually propagate along such long distances? Or do they merely reflect correlations, due to the strong co-expression of neighboring TUs (and/or topoisomerase high-affinity sequences) with shorter-scale propagation of supercoils?

In statistical terms, the first hypothesis would be validated only if TUs of different expression classes were flanked by random sequences and TUs (in terms of distance, orientation, and expression level). This is obviously not true: (1) the mechanism under investigation (transcription-induced supercoils) precisely posits that the expression of neighboring TUs is topologically coupled, and (2) bacterial genomes are the product of evolution, and exhibit synteny, i.e., conserved patterns of neighboring TUs exhibiting strong correlations in expression, in part probably through supercoils (14). Therefore, while the statistical approach carried here does provide a clearer and more quantitative description of the genomic distribution of supercoils (presumably reflecting topological domains), it may only give hints, and no definitive conclusion, regarding the mechanisms underlying them. The reader should therefore be cautious not to over-interpret the observed correlations in terms of causality.

To get further insights into the two hypotheses above, we carried an additional analysis of the binding patterns in the four highest expression classes, by separating each of them depending on the expression level of the neighboring TUs (strongest vs weakest half) in Supplementary Fig. S6. The transcription level in these neighboring regions is considerably weaker than in the central TU (typically ∼100-fold for the strongest class), and for the weakest half, this level is always below the genomic average expression. In spite of this very low expression in flanking regions, the typical pattern of differential topoisomerase binding is present in all analyzed groups, with approximately the same extension (sometimes dropping a little earlier); suggesting indeed that the central TU alone is predominant in the production of supercoils detected in the ∼10 kb range, a lengthscale compatible with previous observations on specific genetic constructions (44). In addition, the groups with more highly expressed neighboring TUs (typically 10-fold higher expression than the weaker ones) also systematically exhibits slightly stronger topoisomerase binding patterns, showing (unsurprisingly) that the latter also contribute to the pattern (especially for topoI, presumably because of the colocalization with RNAP).

Given the density of bacterial genomes, most genes thus presumably share local topological regulatory interactions with at least a dozen of neighbors, presumably resulting in a complex interplay previously hypothesized (14, 43). The coupled expression of neighboring genes thus collectively contributes to the observed strong correlation of expression up to 10 kb apart (13, 14). Analyzing the effect of a single TU would therefore require observing the topoisomerase binding distributions after inducing a specific gene in well-controlled conditions (rather than correlations and averages as we do here). Even in such an experimental setup, we would still expect the expression of neighboring genes to change after the induction, and contribute to the resulting topological landscape.

## Code availability

GRATIOSA is an open-access Python package. It can be automatically installed via PyPI (typically with the command `pip install GRATIOSA`), and is accessible on GitHub with some tutorials, scripts and datasets required to reproduce our analyses: https://github.com/sammeyer2017/GRATIOSA. Documentation is available at: https://gratiosa.readthedocs.io/en/latest/Presentation.html.

