## Supplementary Information for "Quantitative spatial analysis of bacterial transcriptome and chromosome structural data with GRATIOSA: application to twin-supercoiled domain distribution"

Université de Lyon, INSA Lyon, Université Claude Bernard Lyon 1, CNRS
UMR5240, Laboratoire de Microbiologie, Adaptation et Pathogénie, 69621 Villeurbanne, France

**This PDF file includes:**

- Supplementary Figures S1 to S6
- Supplementary Table S1

### Supplementary figures

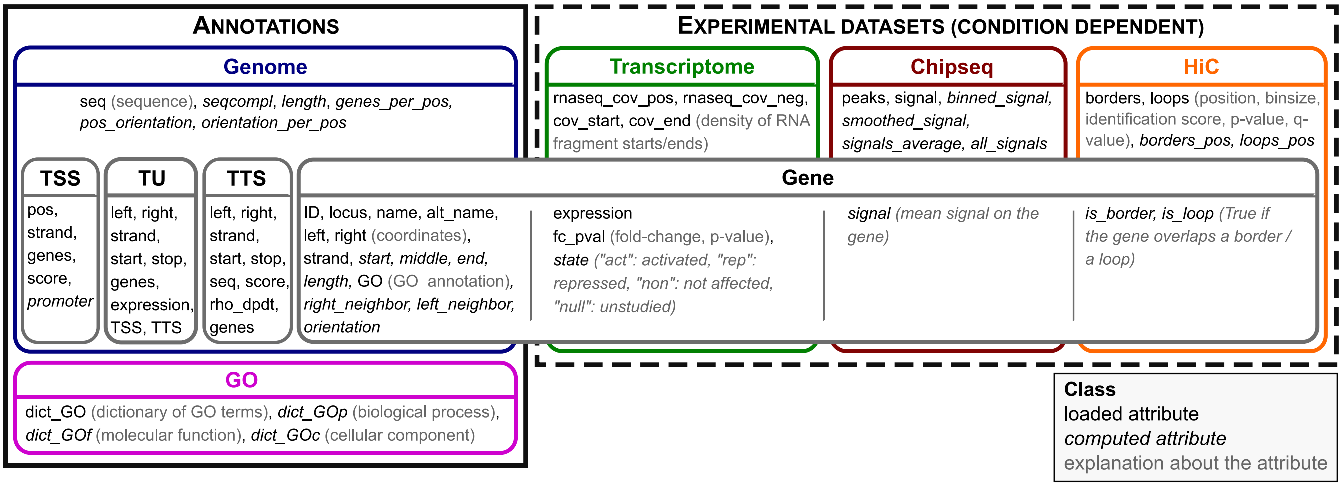


**Supplementary Figure S1**: List of attributes that can be loaded or computed for all classes. The Gene object is an elementary object containing the data of a single gene. Loading the data of all genes is achieved using the Genome objects (for annotations) or either Transcriptome, ChIP-Seq, or Hi-C (for experimental datasets). The same applies to TSS, TU, and TTS objects, whose attributes can be loaded and computed through the Genome class.

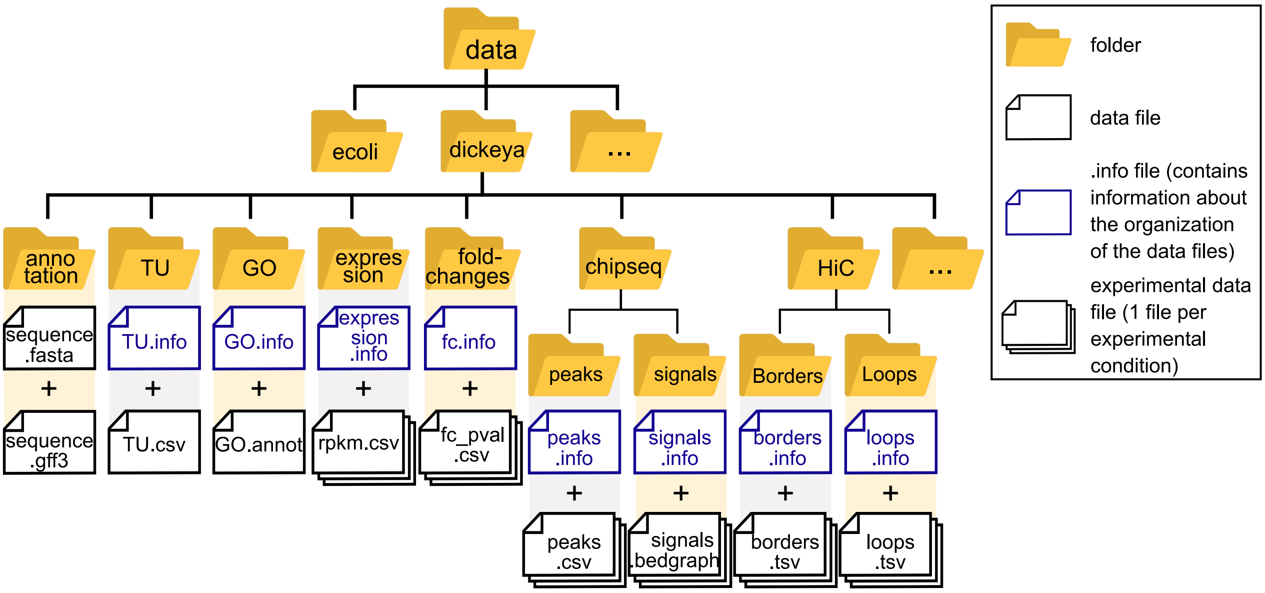


**Supplementary Figure S2**: Structure of the database, containing one directory per organism and one sub-directory for each data type. Each type of experimental data is associated with an info file which allows for the automatic import of a selection of data, of diverse formats.

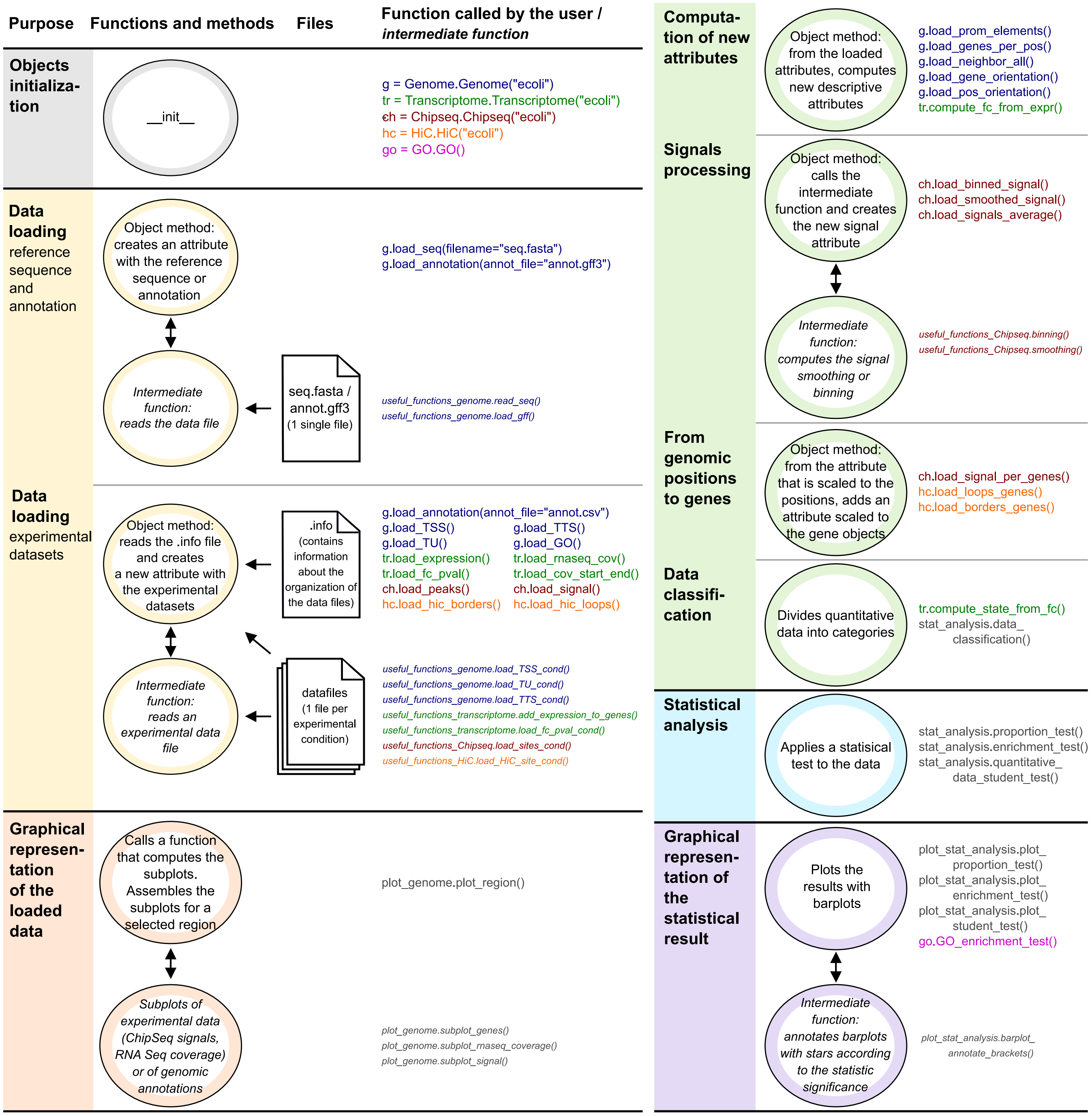


**Supplementary Figure S3**: Standard pipeline for an analysis using the GRATIOSA package. The pipeline consists of four steps (with colors matching those used in Fig 1): objects initialization (gray), data loading as object attributes (yellow), computation of new attributes (green), and statistical analysis (blue). It is also possible to visualize the loaded data (orange) and the statistical results (purple). These steps are performed using the methods associated with each object (Genome, Transcriptome, ChIP-Seq, ...). Each function call is shown in the color of the associated class in Fig S2 (all functions are listed). Function calls written in italics are intermediate functions called within the package, and, in usual cases, not by the user.

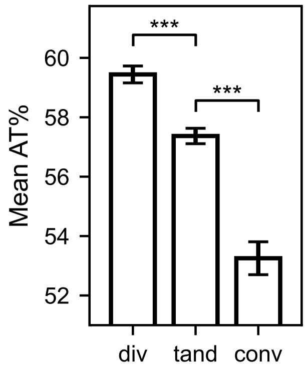


**Supplementary Figure S4**: Average AT content of the intergenic regions by orientation. Error bars are 95% confidence intervals

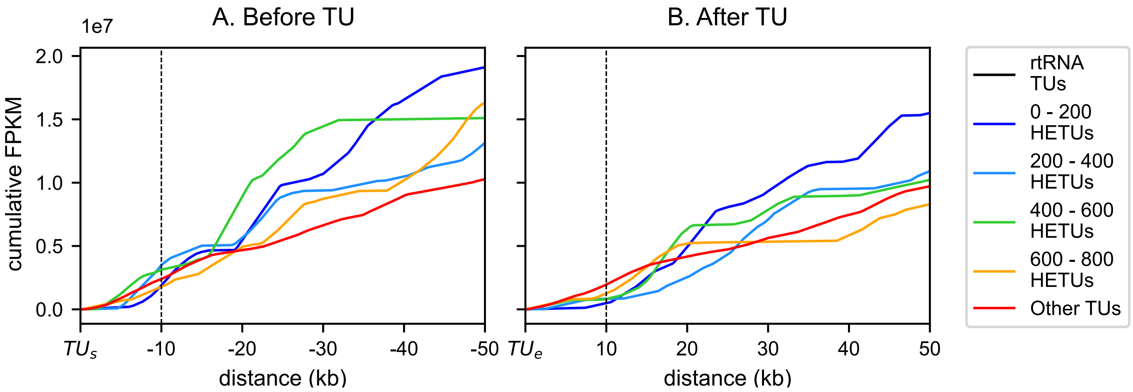


**Supplementary Figure S5**: Cumulative FPKM (expression level) of the neighboring TUs in the 50 kb before the TU start **(A)** and after the TU end **(B)**: the expression of the neighbors does not increase with that of the central TU. TUs were classified based on their expression level, excluding TUs containing stable RNAs (shown in black). The top 200 expressed genes are referred to as 0-200 HETUs, the next 200 most highly expressed genes as 200-400 HETUs, and so on. A FPKM value was associated to each position based on the FPKM values of annotated TUs. If a given position corresponds to multiple TUs, the average FPKM of those TUs is used. The cumulative FPKM represents the sum of FPKM values for each position, from the TU start or TU end to increasing distances (up to 50 kb).

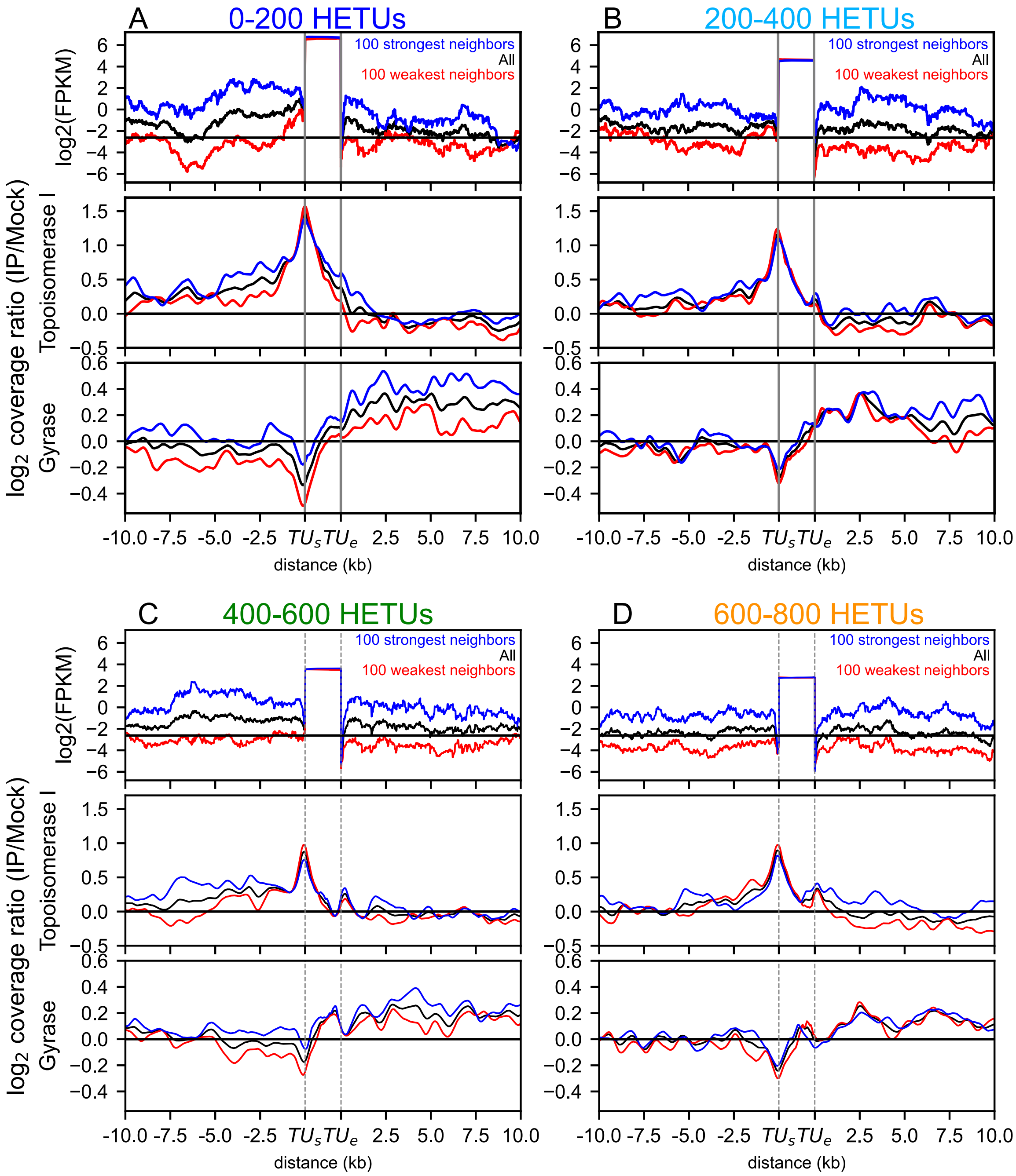


**Supplementary Figure S6**: Role of the neighbors' expression level in the topoisomerase distributions around **(A)** 200 most highly expressed TUs (corresponding to the blue curve in Fig. 2), **(B)** the 200 following (cyan), **(C)** the 200 following (green), **(D)** the 200 following (orange). In each case, the top panel shows the expression level (attributing a homogeneous FPKM level along each TU), the middle panel shows gyrase binding, and the lower panel shows topoisomerase I binding. Black curves shows the average value for all 200 TUs considered, and the blue (resp. red) curve shows the average value for the 100 TUs with highest average expression in the 10-kb flanking regions (versus 100 lowest). In all cases, the central TU has a similar level in the two groups, so that any differences between the red and blue curves are due to the neighbors. For gyrase, a global shift between blue and red curves is sometimes visible (e.g., in A), indicating a larger-scale difference in binding, presumably independent of the expression of the central TU (see text).

#### Supplementary table

**Supplementary Table S1**: Summary table of the statistical tests performed by the package. This table presents the expected inputs, employed modules, and outputs.

| Statistical test | Inputs | Dependencies | Outputs |
|---|---|---|---|
| **Enrichment test** test of the over-representation of a feature in a category | • classification of all items into categories<br>• dictionary containing the feature associated with each item<br>• choice of a targeted feature | • scipy.stats. hypergeom.interval<br>• scipy.stats. hypergeom.sf<br>• statsmodels.stats. multitest. fdrcorrection | For each category:<br>• Number of targeted items (i.e., items associated with the chosen feature) that belong to the category<br>• Expected number of targeted items that should belong to the category according to the category size and the total number of elements having that feature<br>• Proportion of items in the category among all the items associated with the chosen feature.<br>• 95% confidence interval around the proportion<br>• p-value and adjusted p-value of the enrichment test |
| **Proportion test** test of the difference in the proportion of elements associated with a feature between categories | • classification of all items into categories<br>• dictionary containing the feature associated with each item<br>• choice of a targeted feature | • proportion. proportion_confint<br>• statsmodels.stats. proportion. proportions_ztest | For each category:<br>• Proportion of the category items that have the targeted feature<br>• 95% confidence interval around the proportion<br>• P-value of the proportion test (one- or two-tailed test depending on the user's choice) |
| **Student test** test of the difference between the means of different categories | • list of values classified into categories | • scipy.stats .norm.interval<br>• scipy.stats .ttest_ind | For each category:<br>• Mean of the values<br>• 95% confidence interval around the mean<br>• P-value of the student test (one- or two-tailed test depending on the user's choice) |
